# Visualization of the complete primosome reveals the structural mechanisms governing DNA replication restart

**DOI:** 10.64898/2026.01.13.699284

**Authors:** Peter L. Ducos, Alexander T. Duckworth, Kenneth A. Satyshur, James L. Keck, Timothy Grant

## Abstract

Replication restart pathways reinitiate DNA replication processes following their premature termination. In *Escherichia coli*, this essential process begins with the regulated assembly of the primosome complex, comprising the PriA, PriB, and DnaT proteins, onto an abandoned replication fork. Here, we present two distinct primosome structures. One represents an intermediate stage in primosome assembly with a single DnaT C-terminal domain (DnaT^CTD^) bound to PriA/PriB/DNA. The second captures the mature primosome, in which filamentation of multiple DnaT^CTD^ molecules catalyzes the handoff of the single-stranded DNA lagging strand from PriB to DnaT. The DnaT N-terminal domain forms a separate, independent oligomer in the mature structure. Taken together, our results detail the molecular mechanisms underlying replication restart initiation and regulation and suggest an unexpected mechanistic similarity between DnaT and the canonical initiator protein DnaA.

## Introduction

In most bacteria, DNA replication begins at a single origin of replication (*oriC*) within a circular chromosome^1,2^. The initiator protein DnaA binds *oriC* in a sequence-specific manner, leading to oligomerization of the DnaA C-terminal domain (DnaA^CTD^) around DNA and double-stranded (ds) DNA melting^3–5^. The DnaA N-terminal domain (DnaA^NTD^) then self-associates and recruits the helicase/helicase loader (DnaB/DnaC in *Escherichia coli*), and DnaB is loaded onto the newly exposed single-stranded (ss) DNA^3,6–9^. Additional proteins interact with DnaB to form full replication complexes (replisomes)^10^ that drive bidirectional DNA synthesis^11^.

During replication, encounters with damaged DNA, the transcription machinery, or other obstacles can cause the replisome to disengage prematurely, leaving an abandoned replication fork and a partially copied genome^12–15^. Unless the replisome is reloaded, such events are lethal in bacteria^16^. Specialized “replication restart” pathways have evolved to recognize abandoned replication forks in a sequence-independent manner and reload DnaB, reinitiating replication^17,18^. These pathways are also vital for homologous recombination, which generates displacement-loop structures that are also targets of replication restart^18–20^.

In *E. coli*, the major replication restart pathway is driven by the primosome protein complex composed of PriA, PriB, and DnaT. This complex identifies and remodels replication forks, recruits DnaB/DnaC, and directs DnaB loading onto lagging-strand ssDNA to resume DNA replication^17^. PriA initiates the process by binding to DNA in a structure-specific manner^21–25^. PriA is a DNA helicase with two RecA-like domains (PriA^HD1^ and PriA^HD2^) flanked by four additional domains (3′-binding domain, winged-helix domain, cysteine-rich region (PriA^CRR^), and C-terminal domain (PriA^CTD^)) (Figure 1A). PriB, a homodimeric ssDNA-binding protein, is then recruited through direct interactions with PriA and lagging-strand ssDNA^26–28^. The addition of DnaT, which contains N- and C-terminal domains (DnaT^NTD^ and DnaT^CTD^) connected by a linker segment, completes the primosome. The DnaT linker can bind to the ssDNA-binding sites on PriB^29,30^, and the DnaT^CTD^ can form a filament on ssDNA^31,32^, whereas the role of DnaT^NTD^ remains unresolved^33^.

**Fig. 1.**
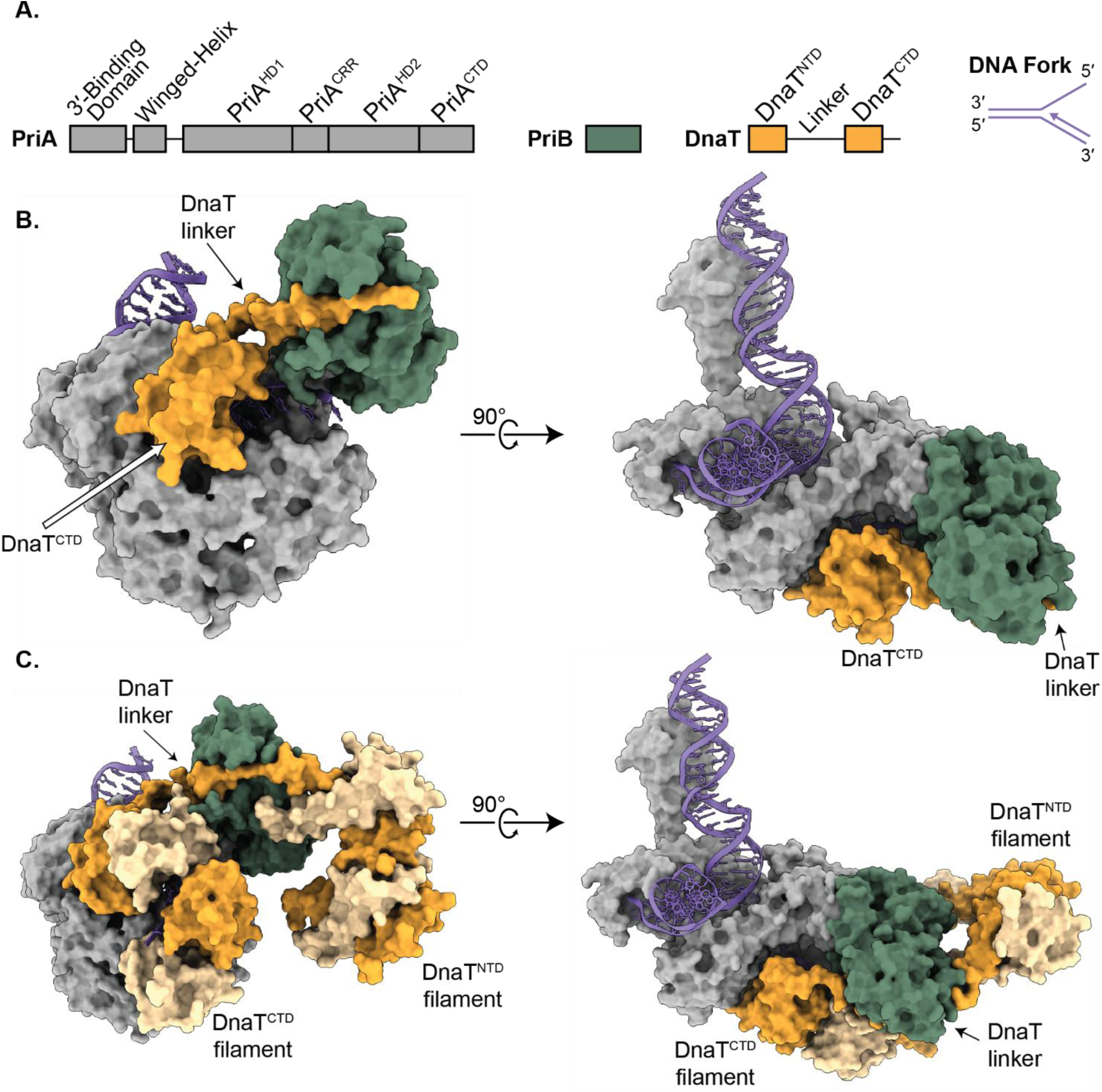
Cryo-EM analysis of the *E. coli* PriA/PriB/DnaT/replication fork complex. **A.** Domain schematics of PriA, PriB, DnaT, and the DNA replication fork. **B and C.** Molecular models of the intermediate **(B)** or mature **(C)** PriA/PriB/DnaT/DNA replication fork complexes. PriA (grey), PriB (green), and DnaT (dark and light orange) are shown in surface representation while the replication fork (purple) is shown as a cartoon.

Our recent cryogenic electron microscopy (cryo-EM) structure of the PriA/PriB/replication fork complex revealed the structural mechanism underlying the first steps of primosome complex formation^34^. PriA recognizes DNA replication forks via multiple contact points with the leading strand, lagging strand, and parental arms of the branched replication fork. The PriA/PriB/DNA structure showed that DNA binding induces a domain rearrangement in which the PriA^CRR^ rotates ∼80° from its position in apo PriA to form a pore that encircles the ssDNA lagging strand. This movement also exposes a surface on PriA^CRR^ onto which PriB docks, positioning PriB to bind lagging-strand ssDNA emerging from the PriA pore.

The structural basis for DnaT addition to the primosome has remained elusive. DnaT is required for DnaB loading onto replication forks *in vitro,* and deletion of the *dnaT* gene in *E. coli* produces severe phenotypes similar to those observed in *priA*-null strains^17,35,36^. Association of DnaT with the PriA/PriB complex requires DNA binding by PriA and is stimulated by PriB^37,38^.

To visualize assembly of the full primosome on replication forks, we analyzed the PriA/PriB/DnaT/replication fork complex via cryo-EM. Two distinct forms of the complex were observed, representing intermediate and mature primosomes. The intermediate structure includes one DnaT^CTD^ and its linker docked onto the replication fork-bound PriA/PriB. The DnaT^CTD^ binds to a surface on PriA formed by the PriA^HD2^ and PriA^CTD^ domains. In prior apo structures, the PriA^CRR^ occludes the DnaT^CTD^ binding site, suggesting that DNA-dependent PriA^CRR^ domain repositioning is also required for DnaT recruitment to the primosome. This parallels the mechanism that regulates PriB binding to PriA^34^. Additionally, the DnaT linker element docks into a known PriB ssDNA-binding groove, likely providing a mechanism for PriB to assist DnaT recruitment to the primosome. The mature primosome structure reveals multiple DnaT^CTD^ molecules forming a helical filament that extends from the single DnaT^CTD^ observed in the intermediate structure. DnaT^CTD^ filamentation is accompanied by a handoff of the ssDNA lagging strand from PriB to DnaT. Additionally, the mature primosome structure contains lower-resolution EM density for a DnaT^NTD^ filament, which is observed projecting away from the remainder of the primosome. Together with previous structural insights, the intermediate and mature primosome structures define a structural model for the PriA/PriB/DnaT DNA replication restart pathway. Intriguingly, the positions of DnaT domains in the mature primosome model reveal an unexpected DnaA-like arrangement in which DnaT^CTD^ filaments on ssDNA to mark the helicase-loading site, and with DnaT^NTD^ element accessible for possible binding to DnaB/DnaC. This parallel suggests that replication restart and canonical replication processes may share significant mechanistic similarities.

## Results and Discussion

### Cryo-EM structures of the *E. coli* PriA/PriB/DnaT/replication fork complex

DNA replication restart in *E. coli* begins with the assembly of a primosome complex, comprising PriA, PriB, and DnaT, on an abandoned DNA replication fork^18,37,38^. Prior individual structures of PriA, PriB, DnaT^CTD^, and of the replication fork-bound PriA/PriB complex have provided important insights into the core primosome proteins and the physical basis underlying the first steps of their assembly into a complex^21,24,26–28,30–32,34^. However, the mature PriA/PriB/DnaT primosome has not been visualized, leaving unanswered questions about the structural mechanisms that regulate primosome formation and the roles of DnaT within the complex. To address this gap, cryo-EM single-particle analysis was used to determine the structure of the complete *E. coli* PriA/PriB/DnaT/replication fork complex.

Purified PriA, PriB, and DnaT (1:2:3 molar ratio, see methods) were combined with a synthetic replication fork consisting of a 25-base pair dsDNA parental arm, a 15-base pair dsDNA leading strand, and a 15-base ssDNA lagging strand (Figure 1A and Table 1). Complexes were isolated using size-exclusion chromatography and frozen onto EM grids. Cryo-EM analysis revealed two major structural classes that yielded high-resolution maps that were used to build two distinct primosome structures (Figure 1, Table 2, and Supplementary Figures 1 and 2). The structures described herein represent ∼50% of the final particles, and each contained PriA, PriB, DnaT, and DNA. Of the remaining particles, 6.8% contained PriA/PriB/DNA fork with no DnaT (representing the previously reported structure^34^), 1.1% contained two PriA molecules that shared a single DNA fork, and the remaining particles classified into lower-quality versions of the reported structures with minor differences, such as small motions of the untethered DNA sections.

**Fig. 2.**
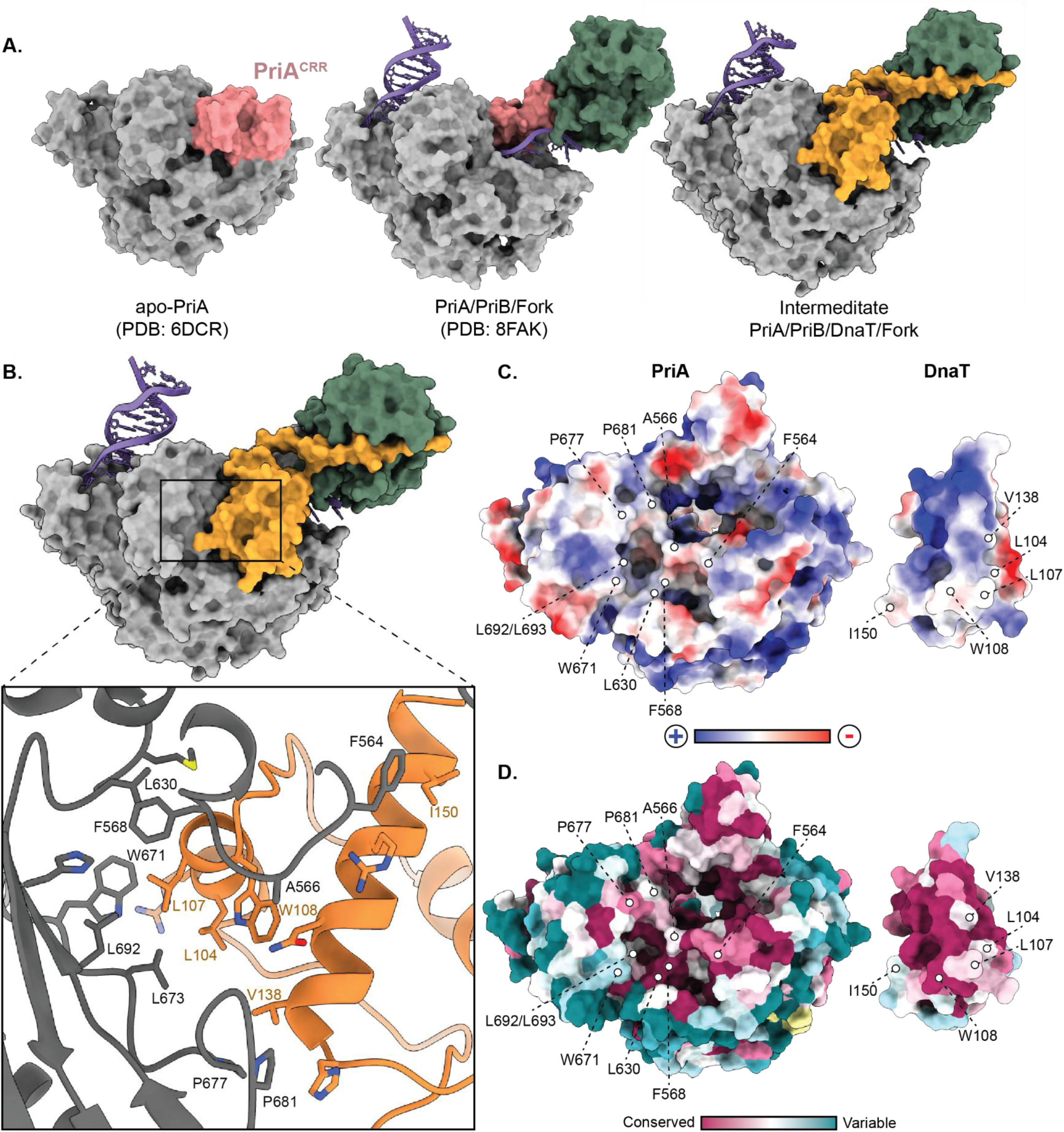
The DnaT^CTD^ interacts with the PriA^CTD^ via a hydrophobic interface. **A.** Models of apo-PriA^24^, the PriA/PriB/replication fork complex^34^, and the intermediate PriA/PriB/DnaT/replication fork complex showing that the PriA^CRR^ (pink) occludes the DnaT (orange) binding site in the apo PriA structure. **B.** Model of the intermediate PriA/PriB/DnaT/replication fork complex containing one DnaT^CTD^ molecule. The inset shows stick model representations of PriA/DnaT residues involved in interaction as assessed by the PISA server^39^. Hydrophobic residues are labeled. **C and D.** Electrostatic potential **(C)** or conservation scores^40^ **(D)** mapped onto the PriA and DnaT binding regions. Key interface residues are highlighted.

**Table 1.**
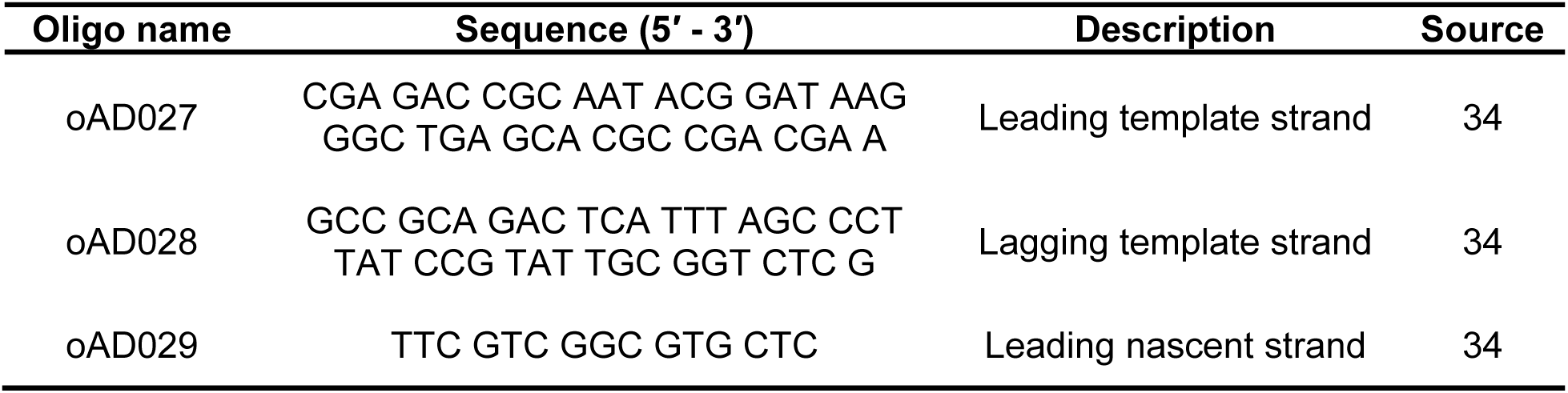
Oligonucleotides used to construct the synthetic replication fork.

**Table 2.** Cryo-EM data acquisition and model statistics.

| **Data collection and processing** | **Intermediate PriA/B/DnaT (EMD-74008, PDB-9ZBT)** | **Mature PriA/B/DnaT (EMD-74009, PDB-9ZBU)** |
| --- | --- | --- |
| Microscope | Krios G3 | Krios G3 |
| Voltage (kV) | 300 | 300 |
| Detector | K3 (Counting) | K3 (Counting) |
| Magnification (nominal/calibrated) | 81,000 | 81,000 |
| Data acquisition software | SerialEM | SerialEM |
| Electron exposure (e<sup>-</sup>/Å<sup>2</sup>) | 100 | 100 |
| Exposure rate (e<sup>-</sup>/pixel/s) | 7.04 | 7.04 |
| Number of frames per micrograph | 102 | 102 |
| Pixel size (Å) | 1.085 | 1.085 |
| Defocus range (µm) | -0.5 to -2.5 | -0.5 to -2.5 |
| Micrographs collected (no.) | 1988 (0° tilt), 1999 (20° tilt) | 1988 (0° tilt), 1999 (20° tilt) |
| **Reconstruction** |  |  |
| Image-processing package | cisTEM | cisTEM |
| Total extracted particles (no.) | 2,200,000 | 2,200,000 |
| Final particles (no.) | 222,000 | 212,000 |
| Symmetry imposed | C1 | C1 |
| Resolution (Å) |  |  |
| FSC 0.143 (unmasked / masked) | 3.7/3.3 | 4.1/3.7 |
| 3DFSC Sphericity | .969 | 0.967 |
| **Model composition** |  |  |
| Protein Residues | 1021 | 1483 |
| Nucleotide Residues | 77 | 80 |
| Ligands | 2 | 2 |
| **Refinement** |  |  |
| Refinement Package | Phenix | Phenix |
| CC (volume/mask) | 0.75/0.74 | 0.73/0.73 |
| Highest resolution used in refinement (Å) | 4.2 | 4.6 |
| R.m.s deviations |  |  |
| Bond lengths (Å) | 0.003 | 0.002 |
| Bond angles (Å) | 0.485 | 0.494 |
| **Validation** |  |  |
| Map-to-model FSC 0.5 | 3.7 | 4.0 |
| Ramachandran plot |  |  |
| Outliers | 0 | 0 |
| Allowed (%) | 1.78 | 2.05 |
| Favored (%) | 98.22 | 97.95 |
| MolProbity Score | 0.95 | 1.10 |
| Poor rotamers (%) | 0 | 0 |
| Clashscore (all atoms) | 1.89 | 2.94 |
| C-beta deviations (%) | 0 | 0 |
| CaBLAM outliers (%) | 1.79 | 1.46 |

The first major structural class (25.04% of particles) yielded a 3.2-Å global resolution map that resolved a PriA monomer, a PriB dimer, a DnaT^CTD^ monomer and linker, and all arms of the DNA replication fork (Figure 1B). This structure represents an intermediate stage in the assembly of the primosome, as only one DnaT molecule is included. The overall architecture of PriA and PriB within the structure was very similar to the previously solved PriA/PriB/replication fork complex structure, with the most notable difference being improved density for the PriA winged-helix domain, which has previously been shown to be flexible relative to the rest of PriA^24,34^. This improvement facilitated rigid-body fitting of the winged-helix domain but was insufficient for detailed molecular modeling. As described in greater detail below, DnaT^CTD^ docks directly onto PriA. Lower resolution density was also observed for the DnaT linker, with none observed for the DnaT^NTD^ in this intermediate structure. The DnaT linker map resolution was insufficient to model the sequence with confidence. However, the map displayed continuous density with the DnaT^CTD^, and it clearly showed the linker bound to a groove on the PriB dimer.

The second major structural class (24.73% of particles) yielded a 3.6-Å global resolution map with density for a PriA monomer, a PriB dimer, and all arms of the DNA replication fork. A DnaT^CTD^ protomer was also included in the same location as that observed in the intermediate structure. The most striking addition in the second structure was the presence of multiple DnaT molecules, including density for a total of four DnaT^CTD^ and DnaT^NTD^ elements, and improved DnaT linker density (Figure 1C). This structure represents the mature primosome.

The DnaT^CTD^ and DnaT^NTD^ oligomers within the mature primosome structure were homotypic, with DnaT^CTD^ and DnaT^NTD^ elements self-interacting and loosely connected by interdomain linker segments. The DnaT^CTD^ oligomer extended away from PriA as a left-handed helical filament. Fourteen of the 15 ssDNA lagging strand bases were resolved in the structure, with eight residing in the DnaT^CTD^ filament interior. The DnaT^NTD^ density was weaker than that for the DnaT^CTD^, but it was sufficient to rigid-body fit two copies of an Alphafold3 predicted dimer structure of the domain^33^. Thus, a four-domain filament of the DnaT^NTD^ was modeled, although the density suggests a longer filament is possible. Continuous density connected one of the DnaT^NTD^ elements to the PriA-bound DnaT^CTD^ via the PriB-bound DnaT linker (Figure 1C).

Weaker density suggested similar connections between the remaining DnaT domains within the structure.

### DnaT docking onto PriA requires DNA-dependent PriA^CRR^ movement

In both primosome structures, a DnaT^CTD^ was observed docked onto a binding site that includes surfaces from PriA^HD2^ and PriA^CTD^ (Figures 1 and 2A). The PriA/DnaT interface buries a surface area of 640 Å^2^ (Figure 2B, calculated via the PISA server^39^). The interface is highly hydrophobic (Figure 2B and C) and is composed of residues that are evolutionarily well conserved (Figure 2D, calculated via the Consurf server^40^, using bacterial species with an annotated *priB* gene). Binding at this position places the DnaT^CTD^ near the PriA pore that encircles lagging-strand ssDNA.

Interestingly, the DnaT-binding site on PriA is not surface-accessible in apo PriA due to the PriA^CRR^ position^34^, suggesting that the PriA^CRR^ must be repositioned for DnaT association (Figure 2A). Relative to its position in apo PriA, the PriA^CRR^ is rotated ∼80° to form the ssDNA pore within the primosome structures. The same structural change was observed in the PriA/PriB/DNA structure^34^, and is needed to expose the PriB binding site on the PriA^CRR^. PriA^CRR^ rearrangement appears to serve as a structural switch that governs access of both PriB and DnaT to PriA, thus regulating primosome formation.

### The DnaT linker docks into a ssDNA-binding groove on PriB

EM density for the linker that connects the DnaT^CTD^ and DnaT^NTD^ was observed in a groove on the surface of PriB in both primosome structures (Figures 1 and 3). The PriB dimer has two basic channels, one in each protomer^26,41^. In the intermediate primosome structure (single-DnaT^CTD^), the lagging-strand ssDNA is bound to the PriB protomer that directly docks onto PriA, consistent with our prior PriA/PriB/DNA structure^34^, whereas the second basic groove is occupied by the DnaT linker (Figure 3A). The DnaT linker density is better defined in the mature primosome structure (oligomeric-DnaT^CTD^) and similarly shows docking into a PriB protomer (Figure 3B). However, as described in greater detail below, the lagging-strand ssDNA is no longer bound to PriB in the mature primosome structure but is instead in the interior of the DnaT filament.

**Fig. 3.**
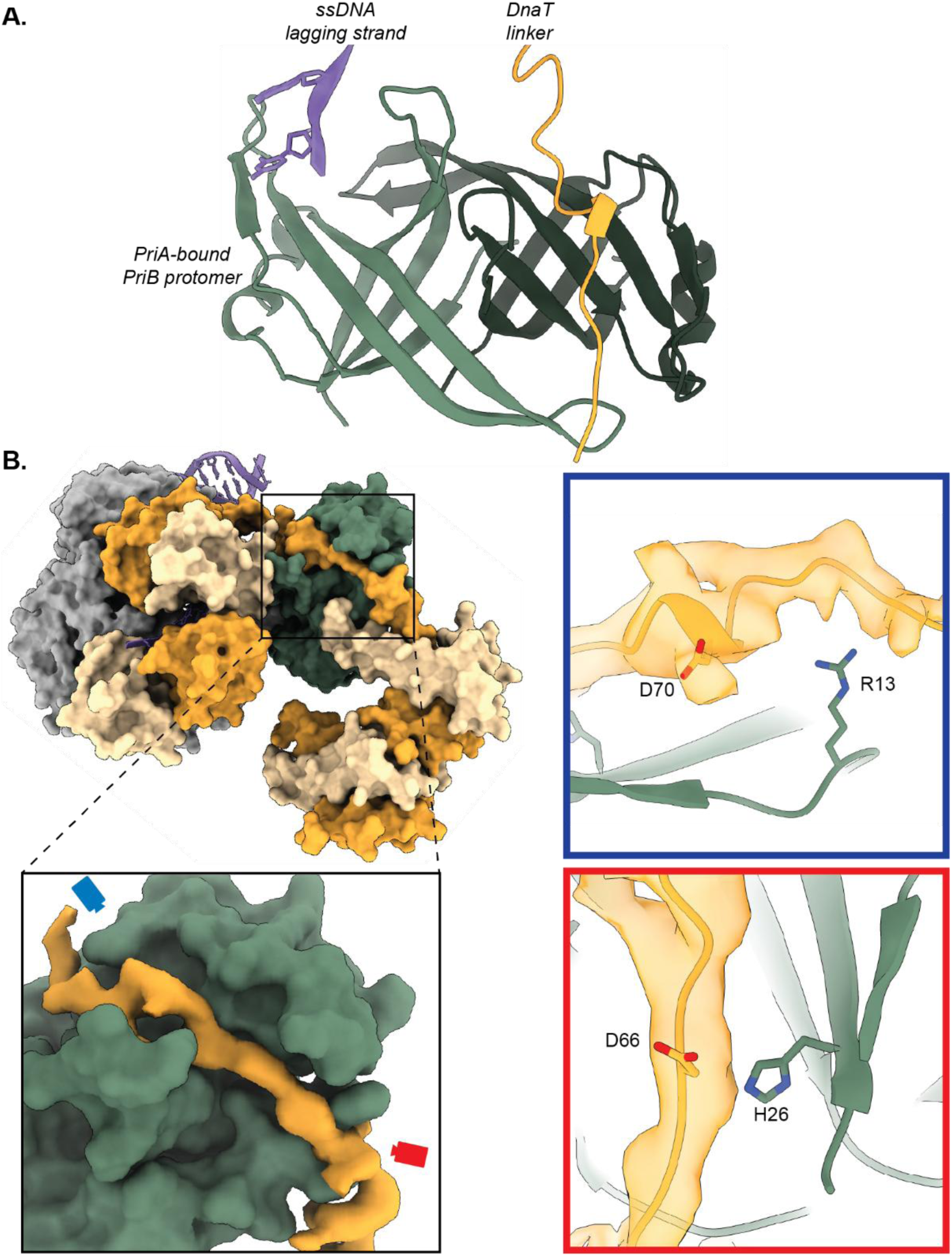
The DnaT linker docks onto a basic groove on PriB. **A.** Model of the intermediate PriA/PriB/DnaT/replication fork complex showing the simultaneous binding of the ssDNA lagging strand (purple) and DnaT linker (orange) to the two PriB protomers (light/dark green). **B.** Model of the mature PriA/PriB/DnaT/replication fork complex containing oligomerized DnaT. The inset shows PriB (green surface) and EM density for the DnaT linker (orange surface). Two views (blue and red boxes) show where the DnaT linker interacts with key PriB residues.

Previous biochemical analysis has shown that PriB facilitates DnaT recruitment to the primosome^37,38^. Two features observed in the structures likely account for this effect: direct interaction between PriB and the DnaT linker and PriB stabilization of the PriA^CRR^ in a position that exposes the DnaT^CTD^ binding site. The DnaT linker is highly acidic, mimicking the negative charge of ssDNA, which facilitates binding to basic grooves on PriB^30,38,42^. PriB residues His26 and Arg13, and DnaT linker residues Asp66 and Asp70 have previously been shown to stabilize the PriB/DnaT interaction^30,38,42^. Due to the inherent heterogeneity of the linker, we could not precisely model the DnaT linker sequence. However, the main-chain model of this region suggests that Asp66 and Asp70 from DnaT are likely near PriB His26 and Arg13, respectively, potentially interacting with one another (Figure 3B).

Although PriB is a homodimer with two potential binding sites for ssDNA or the DnaT linker, the function of this duality has remained unclear. The PriB residues involved in binding the DnaT linker in the primosome partially overlap with those that interact with the PriA^CRR^ and ssDNA. This suggests that PriB is adapted so that one protomer binds to PriA and ssDNA while the second is available to bind the DnaT linker (Figure 3A). This arrangement enables PriA and PriB to coordinate DnaT loading onto the ssDNA, thereby governing primosome maturation so that PriA and PriB binding to a replication fork precedes DnaT association. Such a mechanism could enhance the fidelity of primosome formation and DNA replication restart.

### The DnaT^CTD^ forms a helical filament around lagging-strand ssDNA

As noted above, the cryo-EM map of the mature primosome structure includes density for four DnaT^CTD^ elements. A prior crystal structure of *E. coli* DnaT^CTD^ bound to ssDNA^32^, which formed a helical filament in the crystal lattice, fit the protein density well (Figure 1C). The densities corresponding to the second, third, and fourth DnaT^CTD^ molecules are progressively lower resolution, suggesting flexibility in the filament as it progresses away from the surface of PriA. The DnaT^CTD^ filament surrounds the ssDNA lagging strand (Figure 4). The DnaT^CTD^/ssDNA filament shows similar binding geometry to the previous crystal structure with a nucleotide pair bound between each DnaT^CTD^/DnaT^CTD^ interface (Figure 4C). Because the ssDNA remains bound to PriB when a single DnaT^CTD^ is present in the intermediate complex, our results suggest that DnaT filamentation is required to sequester ssDNA away from PriB. Together with prior biochemical analysis that defined a handoff where DNA is shuttled from PriB to DnaT^29,30,38^, the primosome structures suggest that DnaT linker docking and oligomerization are key for transitioning the lagging strand to DnaT.

**Fig. 4.**
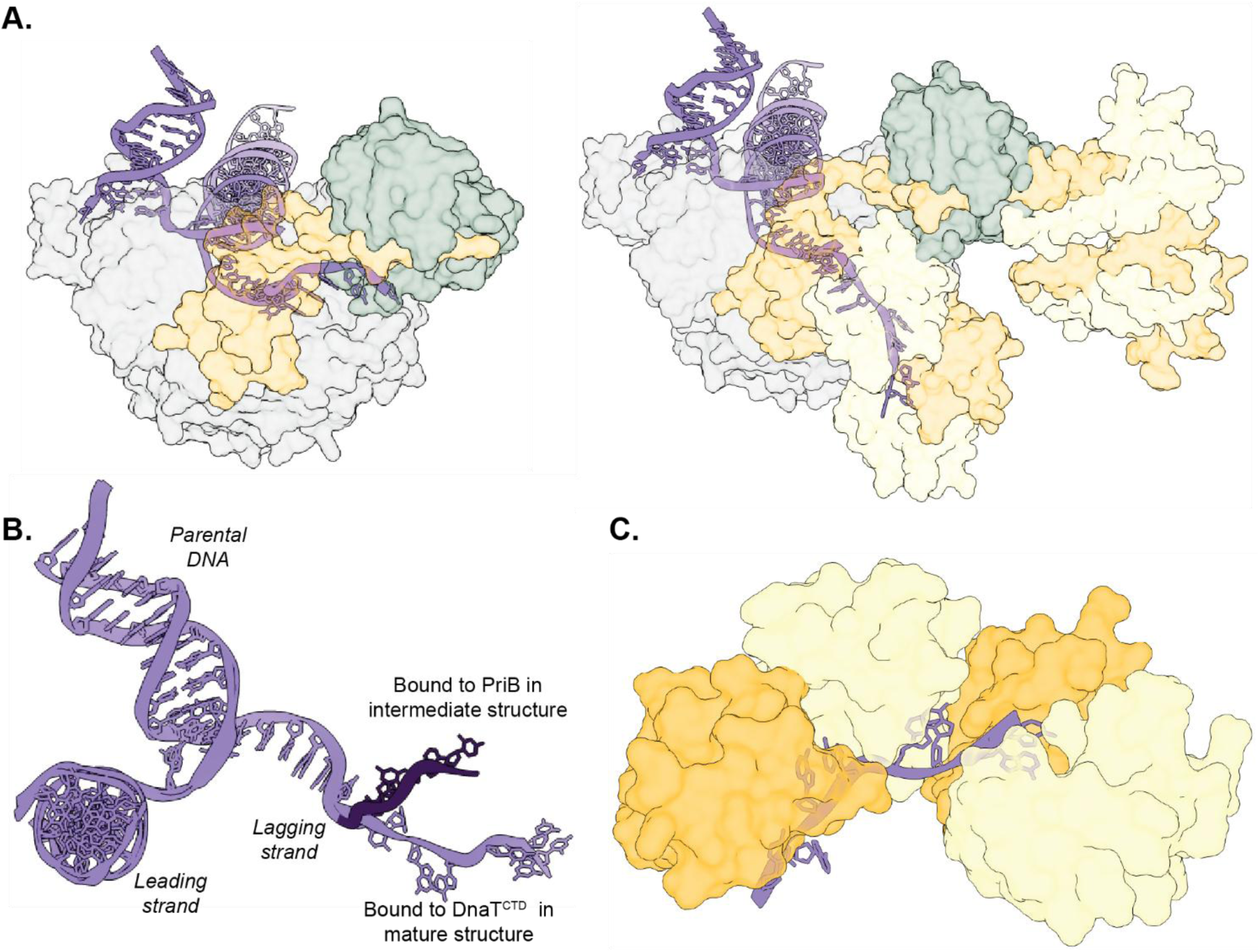
Lagging strand ssDNA is moved from PriB to the DnaT^CTD^ filament. **A.** Transparent models of the intermediate (left) and mature (right) PriA/PriB/DnaT/replication fork structures depicting the different binding orientation of replication fork DNA. **B.** Overlay of the replication forks from both structures. The ssDNA from the intermediate structure is colored dark purple to contrast its divergent path from the mature structure ssDNA. **C.** Close up of the DnaT^CTD^ oligomer bound to DNA showing two bases bound at the junction of each DnaT^CTD^ molecule.

The ssDNA in the mature primosome structure was oriented in the opposite polarity relative to the previous DnaT^CTD^/ssDNA crystal structure^32^. In the primosome, the DnaT^CTD^ filament was aligned in the 3′-5′ direction on the ssDNA, whereas in the crystal structure, the 5′-3′ direction was observed^32^. The reason for this difference is not known, but it could indicate that free DnaT^CTD^ favors ssDNA binding in the 5′-3′ direction. In the context of a replication fork, the directionality of DnaT filamentation is established by its binding to PriA and is thus restricted to 3′-5′ filamentation along the lagging strand ssDNA.

### The DnaT^NTD^ forms a multimer that is stabilized by PriB

Defining the DnaT^NTD^ structure and function has proven elusive. AlphaFold predictions suggest that the DnaT^NTD^ forms dimers, which can further oligomerize into extended filaments^33^.

Consistent with this model, several 2D class averages calculated from the EM data contain filamentous densities spanning large distances, often connecting multiple PriA/PriB assemblies (Figure 5A). Portions of these filaments were also visible in the 3D reconstruction of the mature primosome, although the density was poorly resolved due to conformational flexibility and preferred orientation. Despite this, an AlphaFold3-predicted tetrameric DnaT^NTD^ (comprising two dimers) fit well as a rigid body into the density (Figure 5B)^33,43^. Using the density for the DnaT linker, we connected the first DnaT^CTD^ subunit in the oligomer to a DnaT^NTD^, thereby resolving a full-length DnaT molecule within the complex (Figure 5C).

**Fig. 5.**
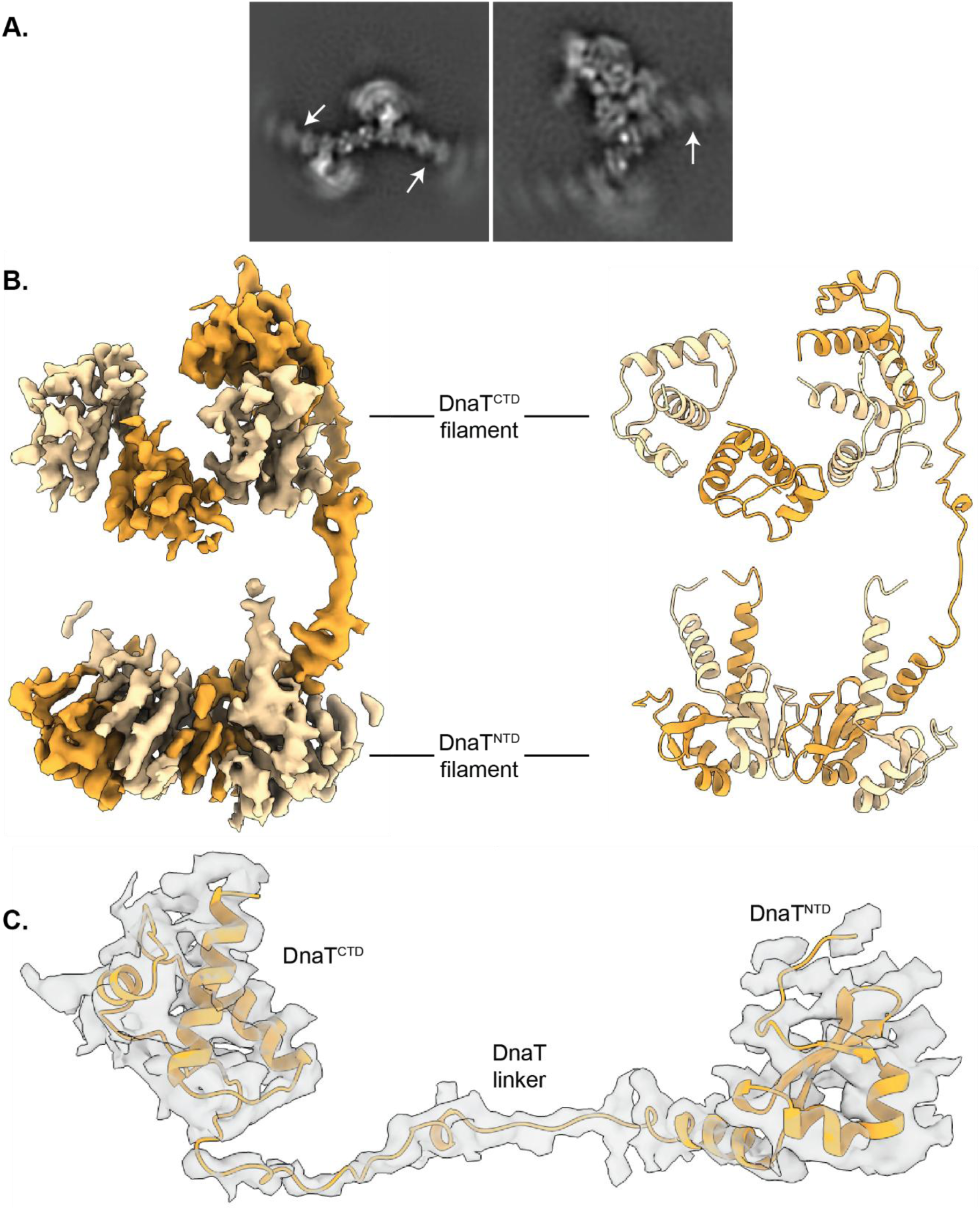
The DnaT^NTD^ forms an extended multimer. **A.** 2D class averages from cryo-EM data processing showing extended DnaT^NTD^ oligomerization (white arrows). **B.** Cryo-EM density (left) and cartoon model (right) of DnaT from the mature PriA/PriB/DnaT/replication fork structure. **C.** Cryo-EM density of a full-length molecule of DnaT from the PriA/PriB/DnaT/replication fork structure.

The resulting structure revealed that the DnaT^CTD^ and DnaT^NTD^ form distinct oligomeric assemblies that coexist within the PriA/PriB/replication fork complex. In addition to its previously described interaction with the PriA^CRR^, PriB is positioned between these two oligomers, establishing contacts with both, as well as with at least one DnaT linker. In this manner, PriB serves as a structural scaffold that supports DnaT filament formation, suggesting a previously unrecognized role for PriB in organizing DnaT architecture during replication restart.

### Structural model of the DNA replication restart pathway

Combined with prior structures of PriA^21,24^, PriB^26–28^, DnaT^CTD 31–33^, and PriA/PriB/DNA^34^, the current structures complete a full description of primosome assembly on a DNA replication fork (Figure 6). The process begins with structure-specific binding to a replication fork by PriA^21–25^. PriA remodels the fork to expose lagging-strand ssDNA, triggering movement of the PriA^CRR^ as PriA encircles the lagging strand. This movement simultaneously exposes the binding sites for PriB (on the PriA^CRR^) and DnaT (on the PriA^HD2^ and PriA^CTD^) to initiate formation of the primosome. PriA^CRR^ movement thus represents the key step linking replication fork recognition to primosome complex formation. PriB docks onto PriA, while also interacting with the lagging strand^26–28^. The primosome structures presented here suggest a stepwise process for DnaT inclusion in the complex with a single DnaT monomer first binding to PriA/PriB/DNA, followed by additional DnaT molecules extending a filament away from PriA. Filamentation of the DnaT^CTD^ is accompanied by the ssDNA lagging strand moving from PriB to DnaT (Figure 4 and 5).

**Fig. 6.**
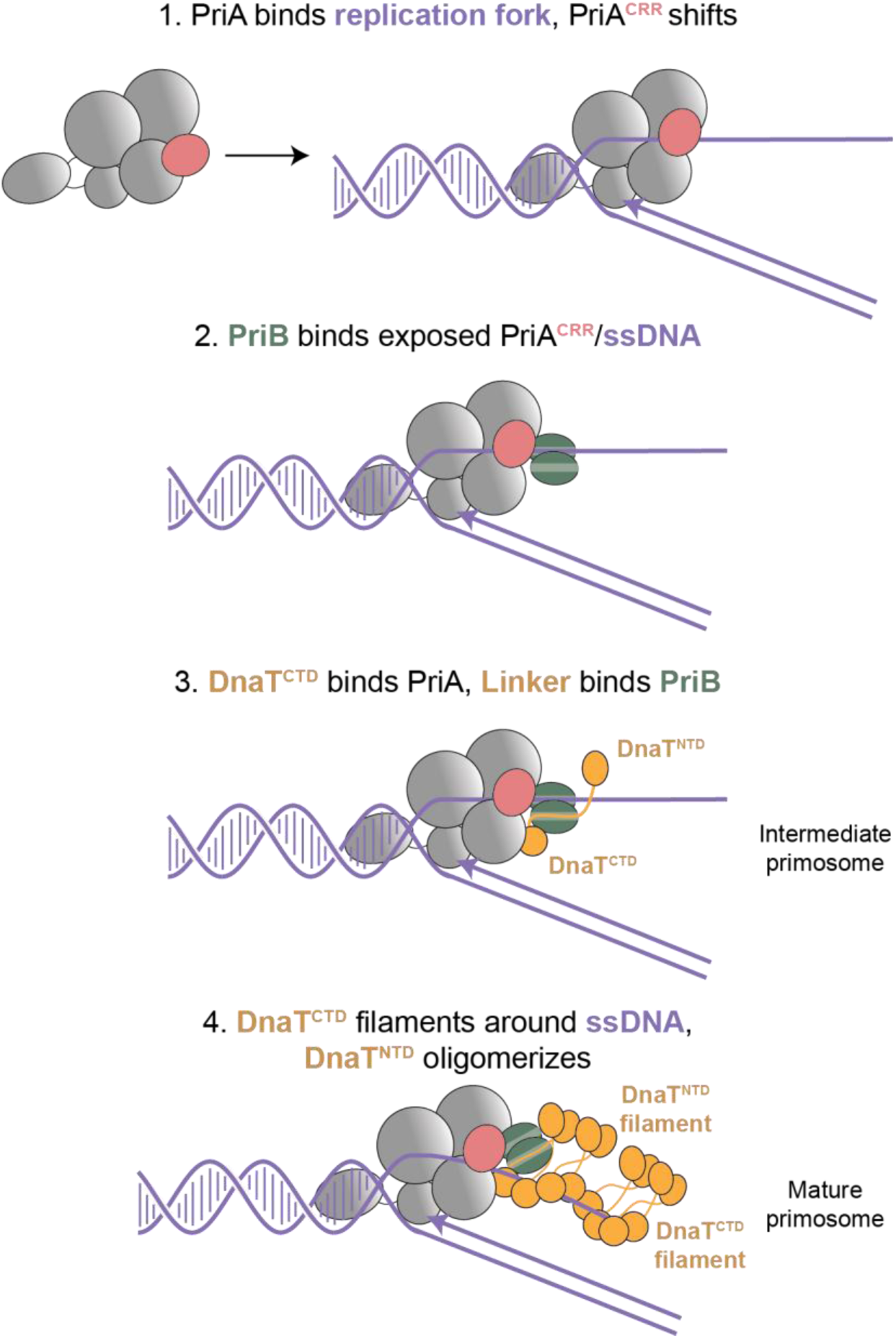
Model of PriA/PriB/DnaT primosome formation. PriA binding to a replication fork triggers rearrangement of the PriA^CRR^ to encircle lagging-strand ssDNA and exposes PriB and DnaT binding sites (1). PriB binds to the exposed binding site of the PriA^CRR^ and ssDNA (2). Once the PriA/PriB/replication fork complex is formed, a molecule of DnaT is recruited through interactions with PriA and PriB (3). Displacement of the ssDNA lagging strand from PriB occurs as additional molecules of DnaT are added, extending a DnaT^CTD^ filament around the ssDNA lagging strand. The DnaT^NTD^ oligomerizes and potentially recruits DnaB/DnaC (4).

### Parallels between DnaT and DnaA suggest a conserved mechanism of helicase/loader recruitment for replication restart and canonical replication initiation

Following primosome assembly, DnaB/DnaC is recruited to the replication fork and a DnaB hexamer is loaded around the lagging-strand ssDNA to reinitiate replication. Although this process has been reconstituted with purified proteins *in vitro*^37,44^, precisely how the primosome recruits DnaB/DnaC remains unknown. The primosome structures presented here have revealed a domain arrangement for DnaT that bears a striking resemblance to that of DnaA. Similarly to DnaT, the DnaA^CTD^ forms filaments on DNA and the DnaA^NTD^ extends away from the DnaA^CTD^/DNA complex^4,5,7–9^. DnaA^NTD^ also forms homodimeric species, which can further associate into high-order oligomeric species that directly interact with DnaB/DnaC^7–9^. DnaA^NTD^ self-association is thought to provide a stabilized binding site for DnaB/DnaC that is not present with monomeric DnaA^9^. The similarities between DnaA and DnaT suggest that DnaT could potentially play analogous roles within the primosome. Consistent with this idea, previous studies have shown genetic interactions between *dnaT* and *dnaC* genes (present in the same operon) that could point to physical interactions between DnaT and DnaC or DnaB/DnaC^35,45,46^.

The DnaA/DnaT parallels could point to general features required to load replicative helicases in both replication initiation and replication restart. The first is a localization factor. For DnaA this is accomplished through sequence-specific DNA binding to *oriC* whereas for replication restart localization is driven by structure-specific DNA binding by PriA. The second is DNA processing to create ssDNA for DnaB loading. For DnaA, processing arises from its intrinsic DNA unwinding activity and filamentation on ssDNA^3,5,8^, whereas for the primosome, processing can include lagging-strand unwinding by the PriA helicase, or movement/exclusion of SSB on the lagging strand by PriA, PriB and DnaT^22,24,47^. Finally, interaction with DnaB/DnaC is required along with DnaB ring loading onto ssDNA. The similar domain arrangements between DnaA and DnaT suggest that DnaT^NTD^ could potentially play a direct role in DnaB/DnaC recruitment. Future experiments are needed to probe parallels between DnaT and DnaA to test this intriguing possibility.

## Acknowledgements

The authors would like to thank the members of the Grant and Keck labs for critical evaluation of this manuscript. We thank the Cryo-EM Research Center (CEMRC) in the Department of Biochemistry at the University of Wisconsin-Madison where the microscopy for this project was performed. Research reported in this publication was supported by National Institutes of Health grant RO1GM098885.

## Methods

### Protein purification

*E. coli* PriA (N-terminal hexahistidine tag) and PriB were expressed and purified as described previously^34,49,50^.

BL21 (DE3) *E. coli* cells transformed with an overexpression plasmid harboring *E. coli* DnaT^38^ were grown shaking at 37 °C in Luria Broth containing 50 μg/mL each of kanamycin and chloramphenicol to an OD_600_ of ∼0.4, then protein expression was induced by the addition of 1 mM isopropyl β-D-1-thiogalactopyranoside for three hours. Cells were pelleted and resuspended in 40 mL Lysis Buffer (50 mM Tris-HCl, pH 8, 10% glycerol, 10% sucrose, 100 mM NaCl, 5 mM ethylenediaminetetraacetic acid, 15 mM spermidine HCl,1 mM phenylmethylsulphonyl fluoride, 1 mM benzamidine, 1 EDTA-free protease inhibitor tablet), lysed by sonication, and clarified by centrifugation. PolyminP was added to the supernatant over 30 minutes to a final concentration of 0.4% (v/v) on ice, the solution was allowed to precipitate for an additional 15 minutes and centrifuged to remove precipitates. Ammonium sulfate was added to the supernatant to a final concentration of 0.163 mg/mL over 30 minutes on ice, protein was allowed to precipitate for an additional 15 minutes and centrifuged to pellet DnaT. The DnaT-containing pellet was resuspended in Resuspension Buffer (10 mM Tris-HCl, pH 8, 10% glycerol, 500 mM NaCl, 2.5 mM ethylenediaminetetraacetic acid) and dialyzed overnight against QFF buffer (10 mM Tris-HCl, pH 8, 10% glycerol, 150 mM NaCl, 2.5 mM ethylenediaminetetraacetic acid). The resulting solution was filtered and loaded onto a HiPrep QFF column (GE) equilibrated with QFF buffer using an ÄKTA Pure FPLC system (GE) at 4 °C. The column was washed and eluted with a linear gradient of QFF buffer (150-1000 mM NaCl). Eluted protein was concentrated and loaded onto a HiPrep S100 column (GE) equilibrated in 575 mM NaCl QFF Buffer. Purified protein was concentrated, dialyzed overnight against storage buffer (10 mM Tris-HCl, pH 8, 50% glycerol, 150 mM NaCl, 2.5 mM ethylenediaminetetraacetic acid), and stored at-20 °C.

### Construction of synthetic DNA replication fork substrates

The synthetic DNA replication fork used in EM was generated as described previously (Table 1)^34^.

### Cryo-EM sample preparation and imaging

PriA (10 nmol), DNA replication fork (10 nmol), PriB (20 nmol, monomers), and DnaT (30 nmol, monomers) were co-incubated in 5 mL S200 Buffer (50 mM Tris-HCl, pH 8, 2 mM dithiothreitol, 5 mM ethylenediaminetetraacetic acid, 75 mM NaCl) on ice. The molar ratio for DnaT was determined under the assumption that it forms a trimer in solution as previously described^51,52^. The sample was centrifuged to remove aggregates and concentrated to 500 μL. The above order of addition prevented aggregation of DnaT. The complex was then purified on a Superdex 200 Increase 10/300 GL analytical size exclusion FPLC column (Cytiva) in S200 Buffer. Peak fractions were combined and concentrated to ∼1 mg/mL. Samples were applied to Ultrafoil 1.2/1.3 grids (Quantifoil) that had been glow discharged for 30 s using a GloQube Plus glow discharge system (Qurom Inc). The grids were plunge frozen at 4 °C using a Vitrobot Mark VI (ThermoFisher) under 100% humidity with a blot time of 4 s.

Movies were collected at 300 kV on a Titan Krios controlled by SerialEM^53^. 1,988 movies were collected using a Gatan K3 camera operating in CDS mode and BioQuantum energy filter with a 20-eV slit width at a calibrated pixel size of 1.085 Å. In a separate session, 1,999 movies were collected using a Gatan K3 camera operating in CDS mode and BioQuantum energy filter with a 20-eV slit width at a calibrated pixel size of 1.085 Å at a 20-degree tilt. The datasets were collected with a total dose of 100 e/Å2 split into 102 fractions. Images were collected with a defocus range of 0.5 to 2.5 μm.

### Cryo-EM data processing

All 3,987 movies were imported into cisTEM 2.065^54^ and processed using the standard workflow. Motion correction and contrast transfer function (CTF) estimation were followed by the exclusion of micrographs with CTF fits worse than 4.0 Å. Approximately 2.2 million particles were automatically picked using the disc picker and subjected to 2D classification, yielding ∼857,000 particles in high-quality classes with clear high-resolution features. These classes showed significant structural heterogeneity. Comparison with previously published 2D class averages^34^ confirmed the consistent presence of PriA and PriB, often accompanied by additional variable densities.

To account for this heterogeneity, an *ab initio* 3D reconstruction was generated from the best 2D classes and refined using auto-refinement with C1 symmetry imposed. An initial 3D classification into five classes was insufficient to resolve the complexity of the sample, so a focused 3D classification was performed with 6 classes focusing the classification around the PriB/DnaT region of the molecule. This enabled the isolation of distinct particle populations. Among these, the map with the best-resolved and highly populated classes corresponding to the PriA/PriB/DnaT/replication fork complex was selected and refined further. The resulting auto-refined 3D volume represented the intermediate primosome structure and accounted for 25.04% of the total particles (∼212,000).

During previous rounds of refinement, diffuse density extending beyond the initial DnaT^CTD^ molecule was present in several classes albeit at very low resolution. To further resolve this density, a 15-class 3D classification was performed on all ∼857,000 particles. One class representing 14.06% of the particles showed more detailed density in this region. To increase the particle count in this class, a 1-round global classification of all particles was performed using this volume as a reference. This led to a volume with greatly improved features in this region. To further improve the detail, a 6-round focused 3D classification was performed focusing on the region determined to be the DnaT filaments. One class representing 24.73% of the particles (∼222,000) showed strong density for the filamentous DnaT regions. This was further auto-refined and resulted in a final 3D volume representing the mature primosome structure. A full overview of the processing workflow is presented in Supplementary Figure 1.

Refinement of the intermediate primosome complex yielded a map with a global resolution of 3.2 Å (FSC 0.143), showing well-resolved density for PriA, PriB, DnaT, and DNA. As anticipated based on their flexibility, the extended DNA arms were unresolved in the final sharpened map. The oligomeric complex refined to a resolution of 3.6 Å (FSC 0.143), clearly resolving PriA, PriB, DnaT, and DNA. The density corresponding to the DnaT^CTD^ oligomer was less well resolved beyond the first subunit in contact with PriA, suggesting conformational heterogeneity. While the remaining DnaT^CTD^ subunits were not sufficiently resolved for *de novo* model building, density for the overall oligomeric assembly remained interpretable.

The map also revealed additional features, including weak but traceable density for the DnaT linker binding to a PriB protomer. In the same oligomeric map, the first helix of the DnaT N-terminal domain was visible, while the rest of the domain and other DnaT^NTD^ subunits were not well resolved, likely due to a combination of preferred orientation and local heterogeneity. To improve the resolution of this region, focused classification was performed, yielding a subset of particles with enhanced density for the linker and proximal DnaT^NTD^, enabling further structural interpretation.

### Model building, refinement, and validation

Model building was carried out using UCSF ChimeraX^55^ and Coot^56^. For the intermediate primosome complex, rigid-body fitting was performed using previously solved structures of the *E. coli* PriA/PriB/replication fork complex (PDB: 8FAK)^34^ and a DnaT^CTD^ unit from the DnaT^CTD^/ssDNA crystal structure (PDB: 4OU7)^32^. For the mature primosome map, the full intermediate model (PDB: 9ZBT) was used as a template, and three additional DnaT^CTD^ subunits with bound ssDNA from the DnaT^CTD^/ssDNA crystal structure (PDB: 4OU7)^32^ were appended by rigid-body docking. For the DnaT^NTD^, an AlphaFold3-predicted^43^ hexameric model of the DnaT^NTD^ was rigid-body fit into the focused classification map. The DnaT linker region connecting the DnaT^CTD^ and DnaT^NTD^ was manually built.

All models were refined using Phenix real-space refinement^57^ and validated using Phenix tools. Refinement statistics are shown in Table 2. Model visualization and figure preparation were performed in ChimeraX and Coot. The final cryo-EM maps for the intermediate and mature PriA/PriB/DnaT/replication fork complexes were deposited in the Electron Microscopy Data Bank (accession codes EMD-74008 and EMD-74009), and the corresponding atomic models were deposited in the Protein Data Bank (accession codes 9ZBT and 9ZBU).

## References

1. Kornberg, A. DNA Replication. J. Biol. CHEMlSTRY 263, 1–4 (1988).

2. Rashid, F. & Berger, J. M. How bacteria initiate DNA replication comes into focus. BioEssays 47, 2400151 (2025).

3. Bramhill, D. & Kornberg, A. Duplex opening by dnaA protein at novel sequences in initiation of replication at the origin of the E. coli chromosome. Cell 52, 743–755 (1988).

4. Erzberger, J. P., Mott, M. L. & Berger, J. M. Structural basis for ATP-dependent DnaA assembly and replication-origin remodeling. Nat. Struct. Mol. Biol. 13, 676–683 (2006).

5. Simmons, L. A., Felczak, M. & Kaguni, J. M. DnaA Protein of *Escherichia coli*: oligomerization at the *E. coli* chromosomal origin is required for initiation and involves specific N-terminal amino acids. Mol. Microbiol. 49, 849–858 (2003).

6. Wahle, E., Lasken, R. S. & Kornberg, A. The dnaB-dnaC replication protein complex of Escherichia coli. J. Biol. Chem. 264, 2469–2475 (1989).

7. Weigel, C. et al. The N-terminus promotes oligomerization of the *Escherichia coli* initiator protein DnaA. Mol. Microbiol. 34, 53–66 (1999).

8. Costa, A., Hood, I. V. & Berger, J. M. Mechanisms for Initiating Cellular DNA Replication. Annu. Rev. Biochem. 82, 25–54 (2013).

9. Abe, Y. et al. Structure and Function of DnaA N-terminal Domains. J. Biol. Chem. 282, 17816–17827 (2007).

10. Lu, Y.-B., Ratnakar, P. V. A. L., Mohanty, B. K. & Bastia, D. Direct physical interaction between DnaG primase and DnaB helicase of *Escherichia coli* is necessary for optimal synthesis of primer RNA. Proc. Natl. Acad. Sci. 93, 12902–12907 (1996).

11. Mok, M. & Marians, K. J. The Escherichia coli preprimosome and DNA B helicase can form replication forks that move at the same rate. J. Biol. Chem. 262, 16644–16654 (1987).

12. Merrikh, H., Machón, C., Grainger, W. H., Grossman, A. D. & Soultanas, P. Co-directional replication–transcription conflicts lead to replication restart. Nature 470, 554–557 (2011).

13. Merrikh, H., Zhang, Y., Grossman, A. D. & Wang, J. D. Replication–transcription conflicts in bacteria. Nat. Rev. Microbiol. 10, 449–458 (2012).

14. Mangiameli, S. M., Merrikh, C. N., Wiggins, P. A. & Merrikh, H. Transcription leads to pervasive replisome instability in bacteria. eLife 6, (2017).

15. Browning, K. R. & Merrikh, H. Replication–Transcription Conflicts: A Perpetual War on the Chromosome. Annu. Rev. Biochem. 93, 21–46 (2024).

16. Cox, M. M. et al. The importance of repairing stalled replication forks. Nature 404, 37–41 (2000).

17. Sandler, S. J. Multiple Genetic Pathways for Restarting DNA Replication Forks in Escherichia coli K-12. Genetics 155, 487–497 (2000).

18. Windgassen, T. A., Wessel, S. R., Bhattacharyya, B. & Keck, J. L. Mechanisms of bacterial DNA replication restart. Nucleic Acids Res. 46, 504–519 (2018).

19. Liu, J. & Marians, K. J. PriA-directed Assembly of a Primosome on D Loop DNA. J. Biol. Chem. 274, 25033–25041 (1999).

20. Masai, H., Asai, T., Kubota, Y., Arai, K. & Kogoma, T. Escherichia coli PriA protein is essential for inducible and constitutive stable DNA replication. EMBO J. 13, 5338–5345 (1994).

21. Bhattacharyya, B. et al. Structural mechanisms of PriA-mediated DNA replication restart. Proc. Natl. Acad. Sci. 111, 1373–1378 (2014).

22. Lee, M. S. & Marians, K. J. Escherichia coli replication factor Y, a component of the primosome, can act as a DNA helicase. Proc. Natl. Acad. Sci. U. S. A. 84, 8345–8349 (1987).

23. Lee, T. Y., Li, Y. C., Lin, M. G., Hsiao, C. D. & Li, H. W. Single-molecule binding characterization of primosomal protein PriA involved in replication restart. Phys. Chem. Chem. Phys. 23, 13745–13751 (2021).

24. Windgassen, T. A., Leroux, M., Satyshur, K. A., Sandler, S. J. & Keck, J. L. Structure-specific DNA replication-fork recognition directs helicase and replication restart activities of the PriA helicase. Proc. Natl. Acad. Sci. 115, 9075–9084 (2018).

25. Windgassen, T. A. & Keck, J. L. An aromatic-rich loop couples DNA binding and ATP hydrolysis in the PriA DNA helicase. Nucleic Acids Res. 44, 9745–9757 (2016).

26. Huang, C. Y., Hsu, C. H., Sun, Y. J., Wu, H. N. & Hsiao, C. D. Complexed crystal structure of replication restart primosome protein PriB reveals a novel single-stranded DNA-binding mode. Nucleic Acids Res. 34, 3878–3886 (2006).

27. Liu, J.-H. et al. Crystal Structure of PriB, a Primosomal DNA Replication Protein of Escherichia coli. J. Biol. Chem. 279, 50465–50471 (2004).

28. Lopper, M., Holton, J. M. & Keck, J. L. Crystal Structure of PriB, a Component of the Escherichia coli Replication Restart Primosome. Structure 12, 1967–1975 (2004).

29. Fujiyama, S. et al. Structure and mechanism of the primosome protein DnaT-functional structures for homotrimerization, dissociation of ssDNA from the PriB·ssDNA complex, and formation of the DnaT·ssDNA complex. FEBS J. 281, 5356–5370 (2014).

30. Fujiyama, S., Abe, Y., Shiroishi, M., Ikeda, Y. & Ueda, T. Insight into the interaction between PriB and DnaT on bacterial DNA replication restart: Significance of the residues on PriB dimer interface and highly acidic region on DnaT. Biochim. Biophys. Acta - Proteins Proteomics 1867, 367–375 (2019).

31. Chen, K.-L., Huang, Y.-H., liao, J.-F., Lee, W.-C. & Huang, C.-Y. Crystal structure of the C-terminal domain of the primosomal DnaT protein: Insights into a new oligomerization mechanism. Biochem. Biophys. Res. Commun. 511, 1–6 (2019).

32. Liu, Z. et al. Crystal structure of DnaT84-153-dT10 ssDNA complex reveals a novel single-stranded DNA binding mode. Nucleic Acids Res. 42, 9470–9483 (2014).

33. Inoue, S., Ikeda, Y., Fujiyama, S., Ueda, T. & Abe, Y. Oligomeric state of the N-terminal domain of DnaT for replication restart in Escherichia coli. Biochim. Biophys. Acta - Proteins Proteomics 1871, 140929–140929 (2023).

34. Duckworth, A. T. et al. Replication fork binding triggers structural changes in the PriA helicase that govern DNA replication restart in E. coli. Nat. Commun. 14, (2023).

35. McCool, J. D., Ford, C. C. & Sandler, S. J. A dnaT Mutant With Phenotypes Similar to Those of a priA2::kan Mutant in Escherichia coli K-12. Genetics 167, 569–578 (2004).

36. Heller, R. C. & Marians, K. J. The Disposition of Nascent Strands at Stalled Replication Forks Dictates the Pathway of Replisome Loading during Restart. Mol. Cell 17, 733–743 (2005).

37. Liu, J., Nurse, P. & Marians, K. J. The ordered assembly of the φX174-type primosome. III. PriB facilitates complex formation between PriA and DnaT. J. Biol. Chem. 271, 15656–15661 (1996).

38. Lopper, M., Boonsombat, R., Sandler, S. J. & Keck, J. L. A Hand-Off Mechanism for Primosome Assembly in Replication Restart. Mol. Cell 26, 781–793 (2007).

39. Krissinel, E. & Henrick, K. Inference of Macromolecular Assemblies from Crystalline State. J. Mol. Biol. 372, 774–797 (2007).

40. Landau, M. et al. ConSurf 2005: the projection of evolutionary conservation scores of residues on protein structures. Nucleic Acids Res. 33, W299–W302 (2005).

41. Szymanski, M. R., Jezewska, M. J. & Bujalowski, W. Interactions of the Escherichia coli Primosomal PriB Protein with the Single-stranded DNA. Stoichiometries, Intrinsic Affinities, Cooperativities, and Base Specificities. J. Mol. Biol. 398, 8–25 (2010).

42. Abe, Y., Ikeda, Y., Fujiyama, S., Kini, R. M. & Ueda, T. A structural model of the PriB–DnaT complex in Escherichia coli replication restart. FEBS Lett. 595, 341–350 (2021).

43. Abramson, J. et al. Accurate structure prediction of biomolecular interactions with AlphaFold 3. Nature 630, 493–500 (2024).

44. Manhart, C. M. & McHenry, C. S. The PriA Replication Restart Protein Blocks Replicase Access Prior to Helicase Assembly and Directs Template Specificity through Its ATPase Activity. J. Biol. Chem. 288, 3989–3999 (2013).

45. Masai, H. & Arai, K. Operon structure of dnaT and dnaC genes essential for normal and stable DNA replication of Escherichia coli chromosome. J. Biol. Chem. 263, 15083–15093 (1988).

46. Lark, C. A., Riazi, J. & Lark, K. G. dnaT, dominant conditional-lethal mutation affecting DNA replication in Escherichia coli. J. Bacteriol. 136, 1008–1017 (1978).

47. Cadman, C. J. & McGlynn, P. PriA helicase and SSB interact physically and functionally. Nucleic Acids Res. 32, 6378–6387 (2004).

48. Tan, Y. Z. et al. Addressing preferred specimen orientation in single-particle cryo-EM through tilting. Nat. Methods 14, 793–796 (2017).

49. Duckworth, A. T., Windgassen, T. A. & Keck, J. L. Examination of the roles of a conserved motif in the PriA helicase in structure-specific DNA unwinding and processivity. PLOS ONE 16, e0255409 (2021).

50. Duckworth, A. T. & Keck, J. L. Use of an unnatural amino acid to map helicase/DNA interfaces via photoactivated crosslinking. in Methods in Enzymology vol. 672 55–74 (Elsevier, 2022).

51. Szymanski, M. R., Jezewska, M. J. & Bujalowski, W. The *Escherichia coli* Primosomal DnaT Protein Exists in Solution as a Monomer–Trimer Equilibrium System. Biochemistry 52, 1845–1857 (2013).

52. Szymanski, M. R., Jezewska, M. J. & Bujalowski, W. Energetics of the *Escherichia coli* DnaT Protein Trimerization Reaction. Biochemistry 52, 1858–1873 (2013).

53. Mastronarde, D. N. Automated electron microscope tomography using robust prediction of specimen movements. J. Struct. Biol. 152, 36–51 (2005).

54. Grant, T., Rohou, A. & Grigorieff, N. cisTEM, user-friendly software for single-particle image processing. eLife 7, e35383 (2018).

55. Pettersen, E. F. et al. UCSF ChimeraX: Structure visualization for researchers, educators, and developers. Protein Sci. 30, 70–82 (2021).

56. Emsley, P., Lohkamp, B., Scott, W. G. & Cowtan, K. Features and development of *Coot*. Acta Crystallogr. D Biol. Crystallogr. 66, 486–501 (2010).

57. Liebschner, D. et al. Macromolecular structure determination using X-rays, neutrons and electrons: recent developments in *Phenix*. Acta Crystallogr. Sect. Struct. Biol. 75, 861–877 (2019).

