## Supplementary Information for "Visualization of the complete primosome reveals the structural mechanisms governing DNA replication restart"

**Affiliations**
[a]John and Jeanne Rowe Center for Research in Virology, Morgridge Institute for Research, Madison, WI 53715, USA.
[b]Department of Biochemistry, University of Wisconsin-Madison, Madison, WI 53706, USA.
[c]Department of Biomolecular Chemistry, University of Wisconsin-Madison, Madison, WI 53706, USA.

[1]Peter L. Ducos and Alexander T. Duckworth contributed equally to this work.

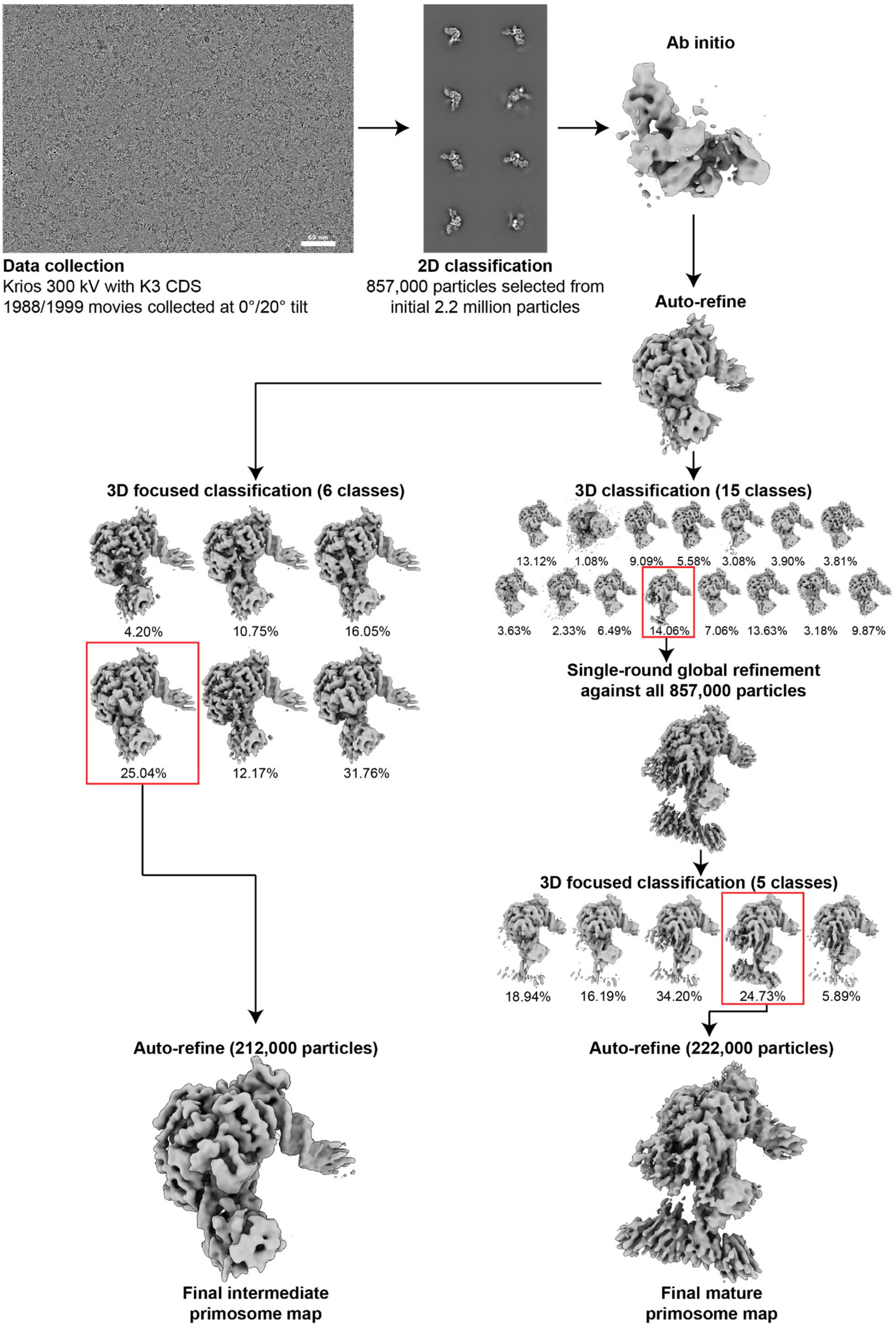
Data collection
Krios 300 kV with K3 CDS
1988/1999 movies collected at 0°/20° tilt
2D classification
857,000 particles selected from
initial 2.2 million particles
Ab initio
Auto-refine
3D focused classification (6 classes)
4.20%
10.75%
16.05%
25.04%
12.17%
31.76%
3D classification (15 classes)
13.12%
1.08%
9.09%
5.58%
3.08%
3.90%
3.81%
3.63%
2.33%
6.49%
14.06%
7.06%
13.63%
3.18%
9.87%
Single-round global refinement
against all 857,000 particles
3D focused classification (5 classes)
18.94%
16.19%
34.20%
24.73%
5.89%
Auto-refine (212,000 particles)
Auto-refine (222,000 particles)
Final intermediate
primosome map
Final mature
primosome map

**Supplementary Fig. 1. Cryo-EM data processing workflow.**
Workflow for cryo-EM structure determination and resulting density maps. Representative cryo-EM micrographs from data collection and selected 2D class averages showing distinct particle views and conformations are shown. 3D reconstructions for all steps in the processing pipeline and particle counts are shown for the intermediate (left) and mature (right) primosome cryo-EM densities.

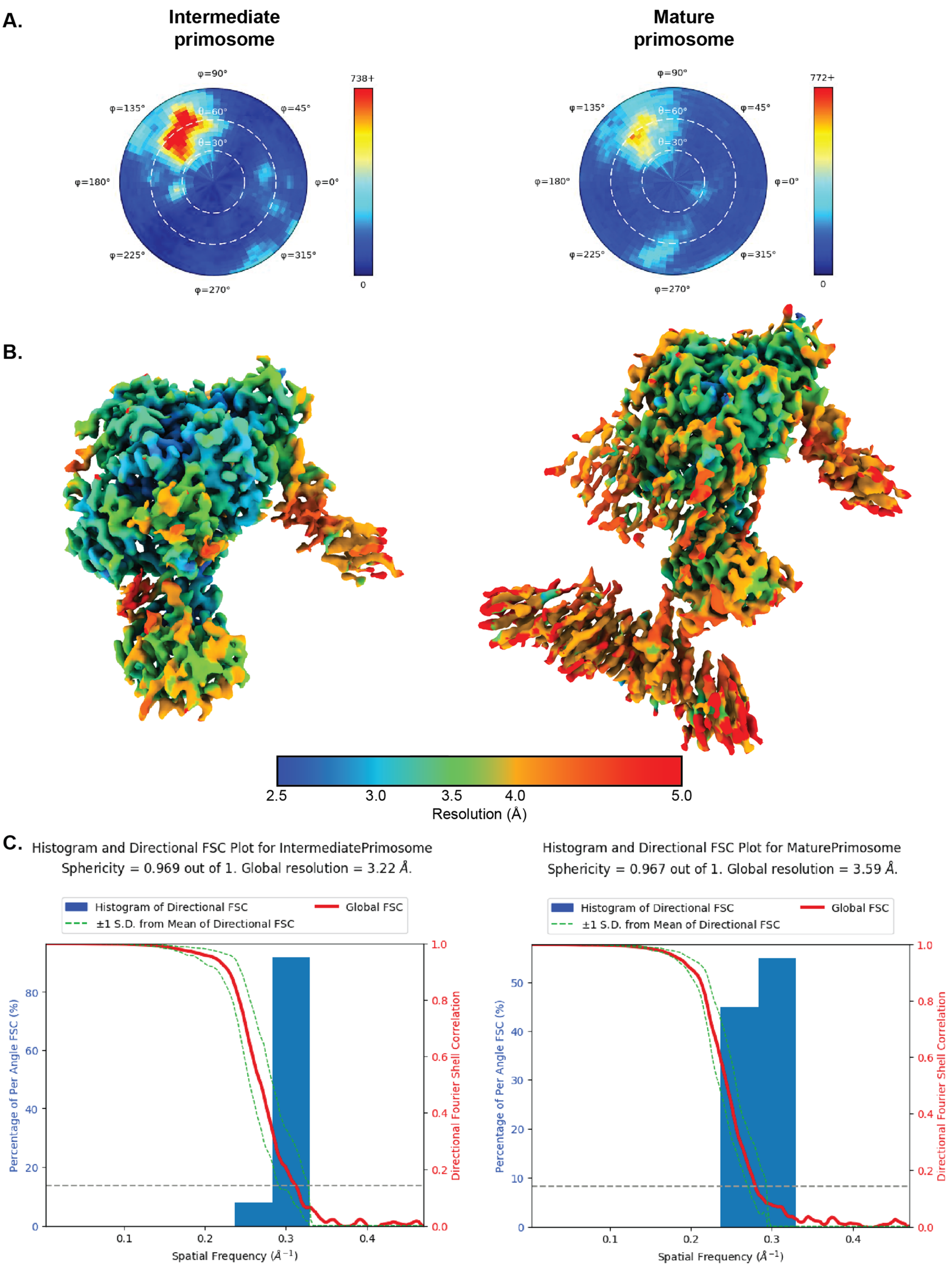
A.
Intermediate primosome
Mature primosome
φ=90°
φ=45°
φ=0°
φ=315°
φ=270°
φ=225°
φ=180°
φ=135°
θ=60°
θ=30°
738+
772+
0
B.
2.5
3.0
3.5
4.0
5.0
Resolution (Å)
C.
Histogram and Directional FSC Plot for IntermediatePrimosome
Sphericity = 0.969 out of 1. Global resolution = 3.22 Å.
Histogram and Directional FSC Plot for MaturePrimosome
Sphericity = 0.967 out of 1. Global resolution = 3.59 Å.
Histogram of Directional FSC
Global FSC
±1 S.D. from Mean of Directional FSC
Percentage of Per Angle FSC (%)
Directional Fourier Shell Correlation
Spatial Frequency (Å$^{-1}$)

**Supplementary Fig. 2. Local resolution and 3D FSC plots.**
**A.** Angular distribution plots illustrating particle orientation coverage for the intermediate (left) and mature (right) primosome cryo-EM densities. **B.** Local resolution maps of the intermediate (left) and mature (right) primosome cryo-EM densities color-coded by resolution from 2.5 to 5 Å. **C.** 3D Fourier Shell Correlation[48] (FSC) plots that illustrate map quality and resolution estimates for the intermediate (left) and mature (right) primosome densities.
